# Genetic detection of RNA-protein interactions using a bacterial three-hybrid assay

**DOI:** 10.64898/2026.06.26.734845

**Authors:** Chandra M. Gravel, Katherine E. Berry

## Abstract

The bacterial three-hybrid (B3H) assay is a powerful genetic tool for detecting interactions between RNA and RNA-binding proteins (RBPs) and assessing the consequences of RBP mutations. This transcription-based system connects the strength of an RNA–protein interaction to the expression of a *lacZ* reporter gene in *Escherichia coli* cells. This *in vivo* approach allows researchers to dissect RNA-protein interactions within a cellular environment, bypassing the need for biochemical purification of RNAs or proteins. This chapter details a three-day protocol for generating quantitative B3H data. Since a significant challenge in B3H assays is RNA misfolding, we describe a recently optimized set of B3H constructs that mitigates this issue by isolating bait RNAs as discrete folding units.

## 1. Introduction

Although RNA-protein interactions are central to post-transcriptional gene regulation, validating these interactions and probing their molecular mechanisms can be technically challenging. The bacterial three-hybrid (B3H) assay presented here [1, 2] has been adapted from the RNA polymerase (RNAP)-recruitment-based principles of an analogous bacterial two-hybrid (B2H) system that detects protein-protein interactions [3, 4]. Together, these assays offer a powerful genetic strategy for studying the molecular interactions of RNA-binding proteins within the *Escherichia coli* cytoplasm. Relative to traditional biochemical approaches such as electrophoretic mobility shift assays (EMSA) or filter binding assays [5, 6], the B3H assay eliminates the need for biochemical purification of each protein or RNA variant. It also serves as a powerful complement to *in vivo* pull-down studies such as RNA immunoprecipitation (RIP)-seq or cross-linking and immunoprecipitation (CLIP)-seq [7–9], by providing a platform for genetic screens to rapidly assess the effects of mutations on the strength of RNA-protein interactions [1, 2, 10]. The B3H assay has successfully detected RNA-protein interactions with K_D_ values up to ∼100 nM, but the limit-of-detection may depend on additional factors beyond binding affinity (see note 1).

The B3H system relies on reporter cells carrying a single-copy *lacZ* reporter gene on an F’ episome. For each experiment, three plasmids are co-transformed to express a tethered tripartite complex consisting of an adapter protein, a prey protein and a bait RNA (Fig. 1) [1, 2]. The pAdapter plasmid expresses a hybrid adapter protein containing the λCI DNA-binding protein fused to the bacteriophage MS2 coat protein (CI-MS2^CP^). This DNA-RNA adapter binds to an O_L_2 operator site upstream of a weak test promoter that drives the expression of a *lacZ* reporter gene. The pBait plasmid directs the arabinose-inducible expression of a hybrid RNA consisting of an MS2 hairpin (MS2^hp^) upstream of a target RNA. The MS2^hp^ anchors the bait RNA to the adapter protein, tethering the target RNA upstream of the test promoter. The pPrey plasmid encodes an IPTG-inducible hybrid protein, containing the N-terminal domain of the alpha subunit (α-NTD) of RNAP fused to the RNA-binding protein (RBP) of interest. If the bait RNA interacts with the prey protein, it stabilizes RNAP at the promoter, activating *lacZ* transcription.

**Figure 1.**
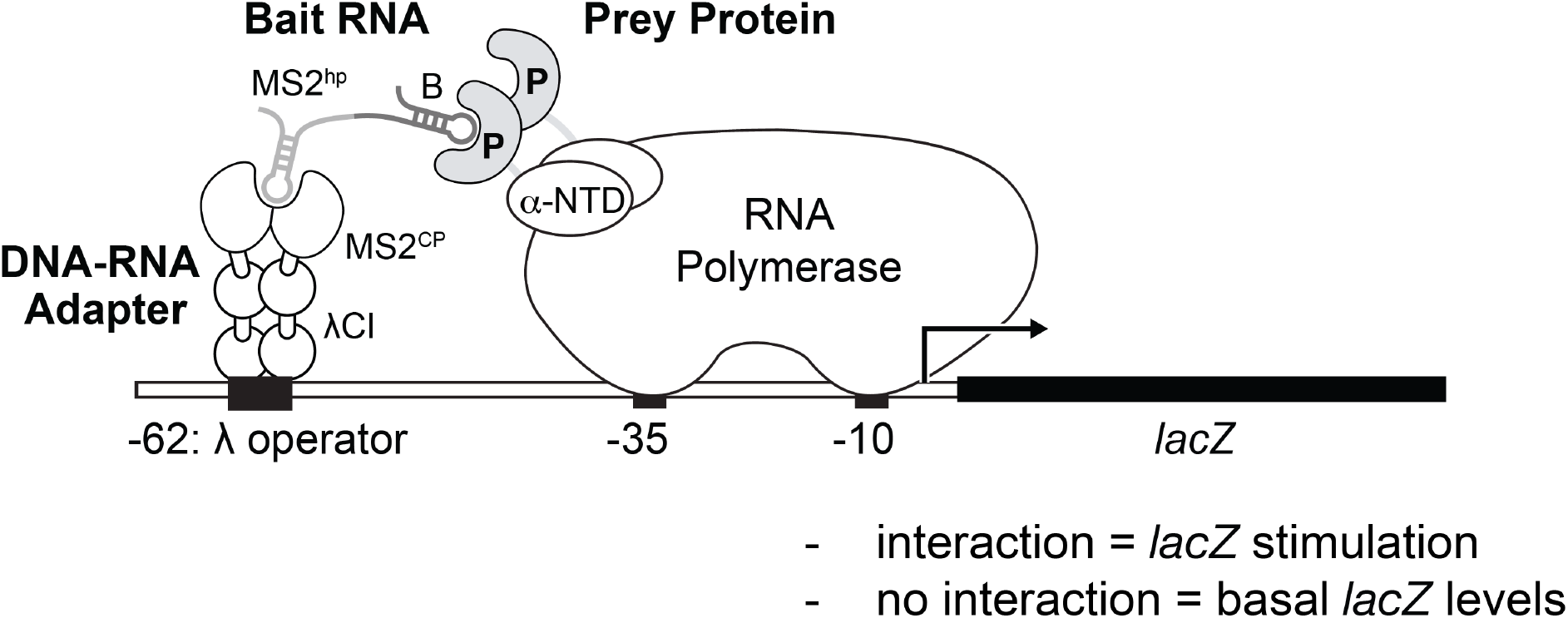
Overview of the bacterial three-hybrid (B3H) system. The bait RNA (B) is fused to a single MS2 RNA hairpin (MS2^hp^), and the prey protein (P) is fused to the N-terminal domain of the α-subunit (α-NTD) of RNA polymerase. The DNA-RNA adapter protein is a fusion of the λCI DNA-binding domain and the MS2 coat protein (MS2^CP^). This adapter protein tethers the bait RNA to a λ operator site located 62 bp upstream of the test promoter. An interaction between the bait RNA and prey protein stabilizes the binding of RNA polymerase to the promoter, activating transcription of the *lacZ* reporter gene, which can be quantified by assaying β-galactosidase activity. **Alternative Text for Figure 1:** A molecular schematic illustrating the architecture of the bacterial three-hybrid system assembled on a segment of DNA. The diagram depicts how three hybrid components interact upstream of a promoter to direct the transcription of a *lacZ* reporter gene. A horizontal line represents the DNA, featuring a lambda operator centered at the minus 62 position, a weak core promoter (with minus 35 and minus 10 elements), and the *lacZ* reporter gene is shown as a solid black box. An arrow pointing into the *lacZ* gene signifies the site of transcription initiation. Bound to the lambda operator is the adapter protein, a fusion of the lambda CI DNA-binding domain and the MS2 coat protein. The MS2 coat protein moiety is shown as a dimer bound to the hybrid bait RNA via an MS2 RNA hairpin. Fused to the MS2 hairpin is the target bait RNA sequence. On the right, a large RNA Polymerase complex is shown. Its alpha-subunit N-terminal domain is fused to the prey protein. The diagram illustrates a positive interaction where the bait RNA physically contacts the prey protein. This interaction acts as a bridge, stabilizing the recruitment of RNA Polymerase to the core promoter. Labels at the bottom of the figure specify that a productive interaction is represented by *lacZ* stimulation, while the absence of an interaction results in basal *lacZ* levels.

The protocol described in this chapter incorporates several recent optimizations over the originally published B3H system that enhance the assay’s signal and reliability. First, the use of a constitutive pAdapter plasmid (e.g., pCW17) prevents previously observed signal inhibition caused by over-expression of an IPTG-inducible adapter protein [11]. Second, updated pBait vectors incorporate three modifications to improve both the versatility of the constructs and the display of target RNAs. These include the use of a single MS2 hairpin (1xMS2^hp^) to minimize the background interaction with a second MS2^hp^ identified with certain RBPs [10]; the introduction of short 5-to 7-bp GC clamps flanking bait RNA sequences to encourage RNAs of interest to fold as discrete structural units [12]; and the addition of an exogenous terminator to facilitate the display of RNAs such as 5′UTRs or open reading frame (ORF) fragments that lack native terminator sequences [12]. These recent iterations of pBait plasmids increased the number of known Hfq-sRNA and -5’UTR interactions that were successfully detected using the B3H assay [12], consistent with them improving the folding and display of bait RNAs.

One strength of this transcription-based B3H system is its compatibility with the classic bacterial two-hybrid (B2H) assay from which it was adapted [3, 4]. Because they share certain components such as the *lacZ* reporter and pPrey plasmids, the two systems can be used together to examine the protein-protein and protein-RNA interactions of a given RBP [13]. While characterizing an RBP’s protein-protein interactions can be interesting for its own sake, these B2H assays can also serve as powerful controls for B3H experiments comparing different variants of an RBP. For example, if a given RBP variant loses its ability to bind RNA in the B3H assay, it can be tested in the B2H system to verify that the prey protein remains properly folded and stably expressed inside the cell. Such an approach can be leveraged in a forward genetic screen [1,2] or serve as an alternative to conducting dot blots or Western blots to assess the stability of particular variants [10].

To implement the B3H system, the protocol below is divided into three stages. First, the target RNA and RBP sequences are amplified and cloned into the optimized pBait and pPrey plasmids. Next, these vectors are co-transformed alongside the pAdapter plasmid into the *E. coli lacZ* reporter strain. Finally, the interaction strength is measured through a β-galactosidase assay.

## 2. Materials

Prepare all solutions using ultrapure water and analytical grade reagents. While general buffers can be stored at room temperature, certain reagents should be stored at specific temperatures required for their stability: IPTG and ONPG stock solutions should be stored at -20°C; antibiotic stock solutions should be maintained at -20°C (or 4°C for kanamycin), and competent reporter cells must be flash-frozen and stored at -80°C to preserve efficiency. In addition, Z-buffer containing β-mercaptoethanol (BME) and ONPG should be prepared fresh for every assay and undiluted BME should be opened in a fume hood.

### 2.1 Plasmids, Strains, and Primers

1. Bacterial cloning strain: *E. coli* NEB 5-α F’I^q^ cells (New England Biolabs) or an equivalent strain providing the *lacI*^*q*^ repressor to ensure repression of IPTG-dependent prey constructs during cloning.
2. Bacterial reporter strains: All B3H reporter strains are derivatives of *E. coli* FW102 [14] carrying a single-copy *lacZ* reporter gene under the control of the *P*_*lac*_ *-62-O*_*L*_2 test promoter on an F’ episome providing kanamycin resistance. FW102 cells contain a streptomycin-resistance gene on the chromosome but selection with this antibiotic is not needed or recommended for B3H assays. Select using kanamycin.
  a. Wild-type strain: FW102 -62-O_L_2 (Addgene #53735; [3])
  b. *Δhfq* strain (KB473; [1]): A derivative of FW102 -62-O_L_2 containing an unmarked chromosomal deletion of the *hfq* gene. Eliminates potential competition by endogenous Hfq for interaction with bait RNAs to enhance interaction signal (see note 2).
3. pAdapter plasmids: Encode the λCI-MS2^CP^ fusion protein, which serves as the DNA-RNA adapter that tethers the hybrid bait RNA to the O_L_2 operator upstream of the test promoter (see note 3). Select using chloramphenicol.
  a. pCW17 (Addgene #174664; [10]): Optimized pAdapter vector that provides a constitutive, lower level of adapter protein and maximizes B3H signal compared to earlier IPTG-dependent variants [11].
  b. pRM16 (Addgene #174787; [2]): Negative control vector corresponding to pCW17; expresses the λCI protein, but lacks the MS2^CP^ moiety.
4. pBait plasmids: Express arabinose-inducible hybrid MS2^hp^ bait RNAs. Select using spectinomycin. See Figure 2 for a schematic overview and notes 4 and 5 for guidance on choosing an appropriate pBait plasmid.
  a. pSS1 (Addgene #222405; [12]): pBait empty control plasmid for RNAs encoding their own transcription terminator; expresses an MS2^hp^ flanked by a 5-bp GC clamp, followed by XmaI and HindIII restriction sites. This plasmid serves as a negative control to define the baseline signal for all 5-bp GC-clamped experimental baits.
  b. pSS2 (Addgene #222406; [12]): pBait empty control plasmid providing an exogenous rho-independent terminator. Expresses an MS2^hp^, followed by XmaI and HindIII restriction sites that are flanked by a 7-bp GC clamp, and a *trpA* terminator (T_*trpA*_) positioned downstream to ensure transcript stability and prevent transcriptional read-through. This plasmid serves as the baseline negative control for baits lacking transcription terminators (*e*.*g*., 5′UTRs).
  c. pLN34 (Addgene #222403; [12]): encodes the *E. coli mcaS* sRNA inserted in the pSS1 backbone; this construct can be digested with XmaI and HindIII to insert new bait RNAs.
  d. pLN43 (Addgene #222404; [12]): encodes the 5′UTR of *E. coli chiP gene* in the pSS2 backbone; this construct can be digested with XmaI and HindIII to insert new bait RNAs.
5. pPrey plasmids: Encodes an IPTG-inducible fusion protein consisting of the N-terminal domain of the α-subunit (α-NTD) of RNA polymerase fused to the RNA-binding protein (RBP) of interest. Select using carbenicillin.
  a. pBr-α (Addgene #53731; [3]): pPrey empty control plasmid expressing the full-length α subunit of RNAP. This construct does not contain NotI and BamHI restriction sites and should not be used to clone prey inserts.
  b. pKB817 (Addgene #174661; [1]): pPrey vector encoding the α-NTD moiety fused to the *E. coli* Hfq protein; this construct can be digested with NotI and BamHI to insert new prey proteins. Interaction between pKB817 (pPrey-Hfq) and pLN34 (pBait-McaS) provides a strong positive control.
6. Sequencing primers: Oligonucleotides that bind to the pBait or pPrey plasmids to allow for F(orward) or R(everse) sequencing of the target insert sequence.
  a. pBait F: 5’-CCGGTAACCCCGCTTATTAAAAGC-3’; binds upstream of the XmaI site on the pBait cloning vectors.
  b. pBait R: 5’-TATCAGACCGCTTCTGCGTTC-3’; binds downstream of the HindIII site on the pBait cloning vectors.
  c. pPrey F: 5’-GAACAGCGTACCGACCTGG-3’; binds upstream of the NotI site on the pPrey cloning vector.
  d. pPrey R: 5’-GGTGATGTCGGCGATATAGG-3’; binds downstream of the BamHI site on the pPrey cloning vector.
7. PCR primers: Oligonucleotides that are complementary to the 5’ and 3’ ends of DNA sequences encoding the bait RNA and prey protein, and also provide appropriate restriction sites. For guidance on designing PCR primers for cloning, see section 3.1.1 and note 6.

**Figure 2.**
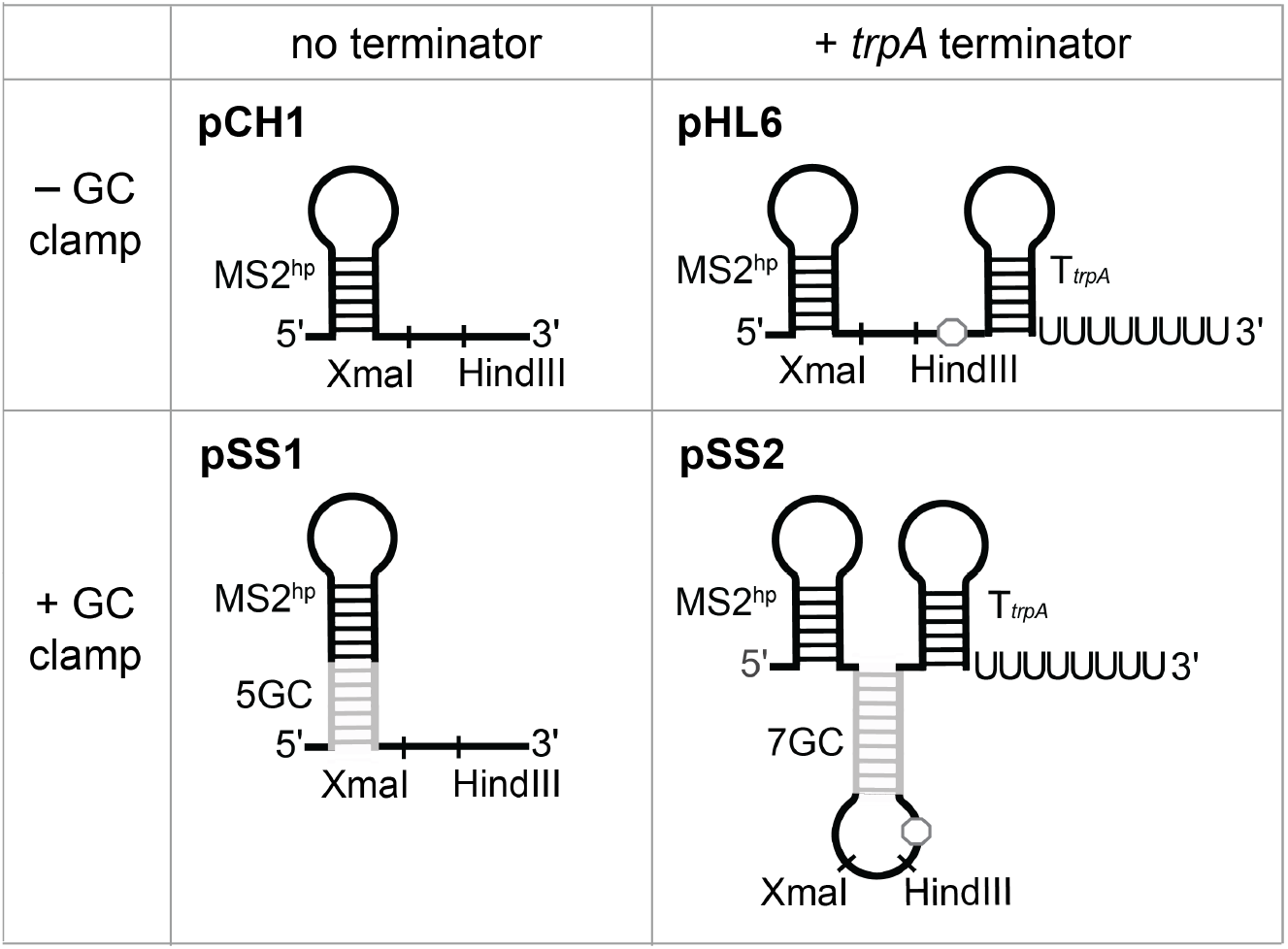
Constructs for display of bait RNA. Schematic highlighting the key structural differences in updated pBait constructs. The simplest pBait construct pCH1 features a single MS2 RNA hairpin (MS2^hp^) followed by XmaI and HindIII restriction sites. pSS1, a derivative of pCH1 containing a 5-bp GC clamp flanking the MS2^hp^ sequence, is generally recommended for displaying bait RNAs that encode their own native rho-independent terminators. The pBait construct pHL6 includes an exogenous rho-independent transcription terminator from the *E. coli trpA* transcript (T_*trpA*_), positioned downstream of the HindIII site. pSS2, a derivative of pHL6 containing a 7-bp GC clamp flanking the bait RNA sequence, is generally recommended for displaying RNAs that lack transcription terminators, such as 5′UTRs or ORF fragments. pHL6 and pSS2 include an UAA stop codon downstream of the HindIII site, represented by a grey octagon (see note 18). For additional information about selecting an appropriate RNA-bait fragment and pBait vector, see notes 4, 5, and 10. **Alternative Text for Figure 2:** A schematic diagram organized into a two-by-two grid comparing four plasmids used for bait RNA display. The horizontal axis distinguishes between constructs with no transcription terminator and those with an exogenous *trpA* terminator. The vertical axis distinguishes between constructs without a GC clamp and those with a GC clamp. Shown in the top left grid is pCH1, represented by a linear RNA sequence starting at the 5-prime end, leading into a single MS2 hairpin stem-loop. This is followed by a short sequence containing the XmaI and HindIII restriction sites at the 3-prime end. The top-right grid shows pHL6. This construct contains the same MS2 hairpin and XmaI and HindIII restriction sites as pCH1 but contains an additional downstream UAA stop codon, represented by a grey octagon, and a second stem-loop followed by an 8 nucleotide poly-uridine tail at the 3-prime end, representing the *trpA* transcription terminator. pSS1 is shown underneath pCH1, in the bottom-left grid. This construct contains the same MS2 hairpin and XmaI and HindIII restriction sites as pCH1, but the base of the MS2 hairpin stem is lengthened with additional grey base pairs and labeled 5GC. pSS2 is shown underneath pHL6, in the bottom-right grid. This is the most complex derivative. Like the other constructs, it features an MS2 hairpin at the 5-prime end. Downstream of this hairpin is a stem loop in which the base-paired region is shown in grey and labeled 7GC. The loop region of this hairpin shows the XmaI and HindIII restriction sites followed by a grey octagon, representing a UAA stop codon. The construct concludes with a trpA terminator sequence as in pHL6 above it.

### 2.2 Media, Reagents

1. Restriction enzymes: NotI-HF, BamHI-HF, XmaI, HindIII-HF, DpnI
2. Antarctic phosphatase
3. T4 DNA ligase
4. High fidelity DNA polymerase (*e*.*g*. Phusion)
5. DNA purification kits: plasmid miniprep kit, gel purification kit, PCR cleanup kit
6. DNA templates: genomic or other template DNA encoding the RNA and protein of interest
7. LB medium: 10 g/L tryptone, 5 g/L yeast extract, 10 g/L NaCl
8. LB Agar: 10 g/L tryptone, 5 g/L yeast extract, 10 g/L NaCl, 15 g/L agar
9. SOC medium: 20 g/L tryptone, 5 g/L yeast extract, 10 mM NaCl, 2.5 mM KCl, 10 mM MgCl_2_, 10 mM MgSO_4_, 20 mM glucose
10. Antibiotic stock solutions: 100 mg/mL carbenicillin in 50% (v/v) ethanol, 25 mg/mL chloramphenicol in 100% ethanol, 50 mg/mL kanamycin in water, 100 mg/mL spectinomycin in water
11. Inducers: 20% (w/v) L-arabinose stock (used at 0.2% final), sterile filtered and stored protected from light, 1M isopropyl-β-D-thiogalactoside (IPTG) stock (used at 0–50 µM final), sterile filtered and stored at -20°C protected from light
12. Z-Buffer: 60 mM Na_2_HPO_4_-7⋅H_2_O, 40 mM NaH_2_PO_4_-H_2_0, 10 mM KCl, 1 mM MgSO_4_, pH 7.0, sterile-filtered
13. ONPG stock solution: 4 mg/mL O-nitrophenyl-β-D-galactopyranoside in Z-buffer. Store in 2 mL aliquots at -20°C.
14. Working assay substrate (Prepare Fresh): Dilute 4 mL of ONPG stock into 16 mL of Z-buffer and add 43.2 μL β-mercaptoethanol (BME) (final concentration: 0.8 mg/mL ONPG, 0.22% (v/v) BME (see note 7).
15. PopCulture reagent (Novagen/Millipore Sigma)
16. rLysozyme (Novagen/Millipore Sigma). Dilute to 400U/µL in its storage buffer (100 mM NaCl, 50 mM Tris-HCl, 1 mM DTT, 0.1 mm EDTA, 50% glycerol, 0.1% Triton X-100, pH 7.5) and store at -20°C.
17. Working lysis buffer: Add 1.2 μL rLysozyme (400U/µL) to 1.2 mL of PopCulture Reagent. This is enough to fill one 96-well plate.
18. Agarose for DNA gels

### 2.3 Equipment

1. Microplate spectrophotometer: Capable of reading absorbance at 420nm and 600nm (OD_420_ and OD_600_; *e*.*g*., Molecular Devices SpectraMax 190)
2. Microplate shaker: Capable of shaking at 900 rpm and housed within a 37°C incubator
3. PCR thermocycler
4. 96-well deep-well (2 mL) blocks
5. Gas-permeable breathable film
6. 96-well flat-bottom clear polystyrene plates with lids for 96-well bacterial growth (*e*.*g*., Greiner Bio-One)
7. Multichannel pipettes: 8- or 12-channel pipettes (P10, P200)

## 3. Methods

The B3H assay is performed in three major stages, each detailed below: molecular cloning (3.1 and 3.2), reporter-cell transformation (3.3), and interaction measurement (3.4; see notes 8 and 9 for possible variations). Figure 3 provides a schematic overview of steps 3.2-3.4.

**Figure 3.**
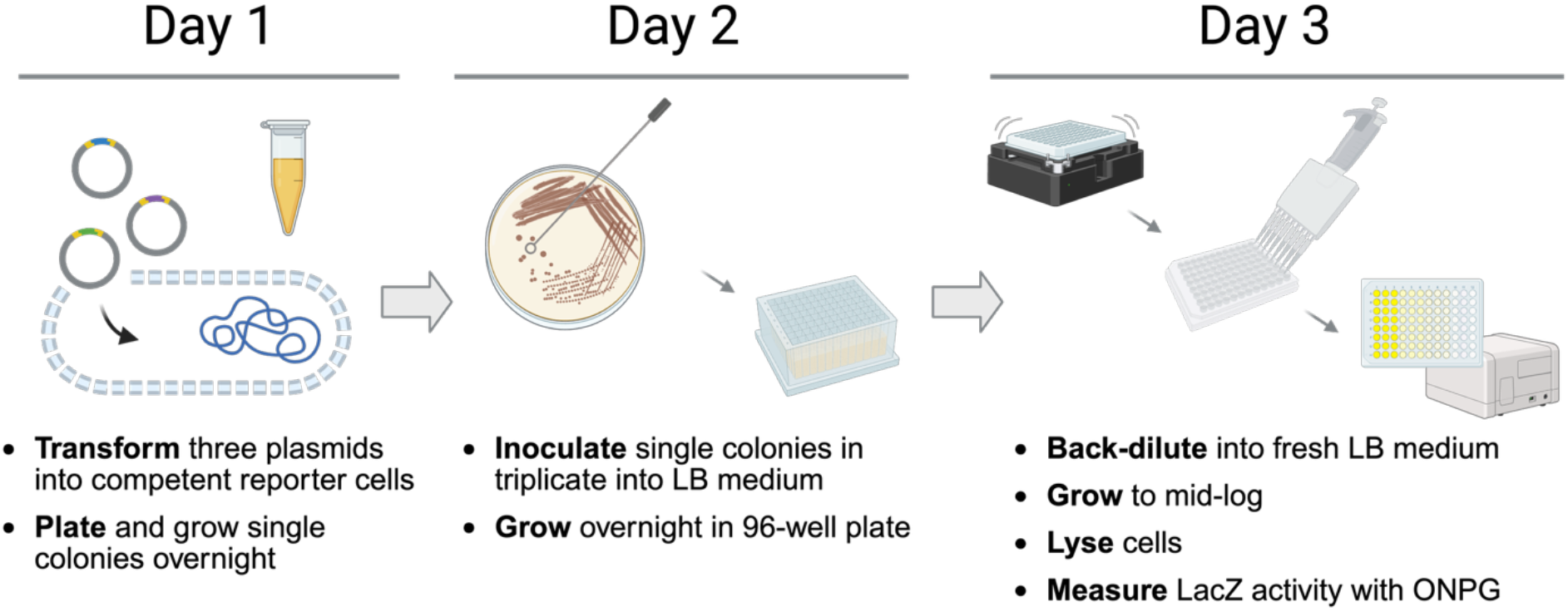
Overview of the workflow for a three-day B3H protocol. On day one, plasmids pPrey, pBait, and pAdapter are co-transformed into competent *lacZ* reporter cells and grown on LB agar plates supplemented with four antibiotics. On day two, single colonies are inoculated into liquid LB supplemented with inducers and antibiotics in a 96-well deep-well plate. On the third day, overnight cultures are back-diluted into fresh LB medium supplemented with antibiotics, L-arabinose and IPTG. After growing to mid-log phase, cells are lysed and β-galactosidase activity is measured via the rate of ONPG turnover, detected spectroscopically. See sections 3.3 and 3.4 for detailed protocols. Figure made using Biorender.com. **Alternative Text for Figure 3:** A workflow diagram divided into three sequential panels, illustrating the progression of the three-day bacterial three-hybrid (B3H) protocol. The first panel, labeled Day 1, shows a schematic of an eppendorf tube, with three circular plasmids being introduced into a dashed oval representing a competent *lacZ* reporter cell. Text under this panel reads “Transform three plasmids into competent reporter cells; Plate and grow single colonies overnight.” The second panel, labeled Day 2, depicts a sterile toothpick picking a single colony from an agar plate with an arrow pointing to a 96-well deep-well block. Text under this panel reads “Inoculate single colonies in triplicate into LB medium; Grow overnight in 96-well plate.” The final panel is labeled Day 3. A 96-well plate is shown on a microplate shaker, with an arrow pointing to an image of a multichannel pipette adding liquid to a flat-bottom 96-well plate. An additional arrow points to a 96-well plate with varying intensities of yellow color next to a microplate spectrophotometer. Text under this panel reads “Back-dilute into fresh LB medium; Grow to mid-log; Lyse cells; Measure *LacZ* activity with ONPG.”

### 3.1 Primer Design and Insert Preparation

1. Design primers to amplify the desired protein and RNA of interest and introduce appropriate restriction sites for vector ligation (see notes 6 and 10).
  a. For RNA baits, include an XmaI site on the forward primer and a HindIII site on the reverse primer.
  b. For protein preys, include a NotI site on the forward primer and a BamHI site on the reverse primer.
  c. To ensure efficient restriction digestion of the PCR products, include 4–8 nt ‘landing pads’ at the 5′ ends of all primers.
  d. Crucially, for prey constructs, an extra nucleotide (typically an ‘A’) must be included immediately following the NotI site in the forward primer to maintain the proper reading frame with the α-NTD.
  e. If the sequence encoding the prey protein lacks its own stop codon, introduce one upstream of the BamHI site in the corresponding reverse primer.
2. Amplify the DNA sequence encoding the RNA or protein of interest via PCR using a high-fidelity DNA polymerase and template DNA. If a plasmid with a matching antibiotic resistance gene was used as the PCR template, add 1 µL of DpnI to the reaction and incubate for 1 hour at 37°C.
3. Verify the PCR product by resolving 5 µL of the PCR reaction on an 1% agarose gel to confirm a single band of the expected length.
4. Isolate the remaining PCR product with a PCR cleanup kit.
5. Digest the PCR product with the appropriate restriction enzymes (1 µL each). Incubate for 1-2 hours at 37°C.
6. Purify the digested insert using a PCR cleanup kit.

### 3.2 Vector Preparation and Cloning

1. Select an appropriate pBait vector to display your RNA of interest (Fig. 2), based on whether it encodes its own transcription terminator (see note 4). The use of constructs with GC clamps is generally recommended to promote correct folding of bait RNAs (see note 5).
2. Digest 2–5 µg of the appropriate B3H vector (*e*.*g*. pSS1-derived pLN34 or pSS2-derived pLN43 for pBait and pKB817 for pPrey) with the corresponding restriction enzymes. Incubate for at least 2 hours and up to overnight at 37°C (see note 11).
3. Add 1µL antarctic phosphatase directly to the digestion reaction and incubate 1h at 37°C to dephosphorylate the 5’ ends and prevent self-ligation.
4. Gel purify the digested vector on an 1% agarose gel to separate from any undigested parent plasmid.
5. Ligate digested vector and insert at a 1:3 molar ratio using T4 DNA Ligase and incubate 1h at room temperature or overnight at 4°C.
6. Transform the ligation mixture into a *lacI*^*q*^ *E. coli* cloning strain. Plate on selective LB agar (carbenicillin for pPrey, spectinomycin for pBait) and incubate overnight at 37°C.
7. Identify correct clones by plasmid miniprep and Sanger sequencing (see section 2.1.6 for appropriate sequencing primers).

### 3.3 Transformation and Culture Growth

1. To prepare for transformations, pipette 1µL of each of the three plasmids (pPrey, pBait, pAdapter) into 30 µL chemically competent FW102 -62-O_L_2-*lacZ* reporter cells (see note 12). Along with the experimental transformation containing all three hybrid components (pPrey + pBait + pAdapter), include three negative control transformations, each lacking one of the three hybrid constructs: (1) pPrey(α)-empty + pBait + pAdapter, (2) pPrey + pBait(MS2^hp^)-empty + pAdapter and (3) pPrey + pBait + pAdapter-empty (Fig. 4; see notes 3, 13, and 14).
2. Let cells and plasmids incubate on ice for at least 20 minutes, then heat shock for 45 seconds at 42°C and place back on ice for 2 minutes.
3. Add 500 µL SOC medium and allow cells to recover 1 hour at 37°C with gentle shaking.
4. Plate transformations on LB agar plates containing 4 antibiotics (carbenicillin, chloramphenicol, spectinomycin and kanamycin; these select for the three plasmids and reporter strain, respectively) and grow overnight at 37°C.
5. Inoculate single colonies from transformation plates into 1 mL of LB medium containing the same 4 antibiotics and 0.2% L-arabinose in a 96-well deep-well plate. IPTG addition at this stage is optional (see note 15). Inoculate at least three independent colonies per transformation to serve as biological replicates.
6. Cover this deep-well plate with a gas-permeable breathable film and incubate at 37°C with vigorous shaking (approx. 900 rpm) overnight (∼12–16 hours).
7. Subculture 5 µL of the overnight growth into 200 µL fresh medium supplemented with the same antibiotics and arabinose in optically clear 200 µL flat bottom 96-well plate. The concentration of IPTG in these subcultures can be varied (see note 15). Cover plate with a plastic lid and grow cells to mid-log phase (OD_600_ approx 0.4–0.8) at 37°C with vigorous shaking (see note 16).

**Figure 4.**
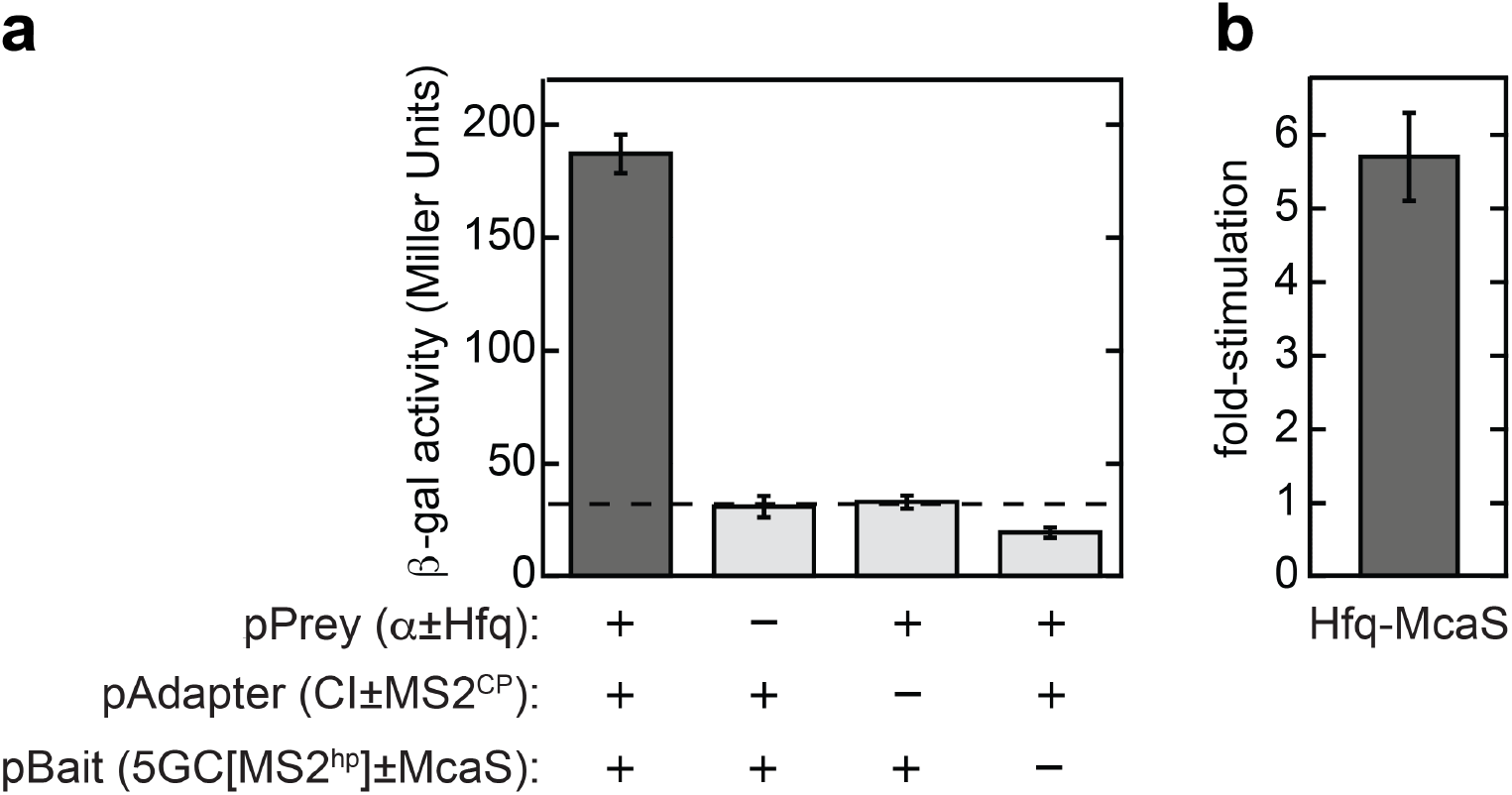
Example B3H data of an Hfq-RNA interaction. Representative data for B3H interaction of *E. coli* Hfq and McaS. (a) Raw β-galactosidase levels of reporter cells (KB473) transformed with pKB817 (pPrey-Hfq), pCW17 (pAdapter) and pLN34 (pBait-McaS) or corresponding negative controls in which one hybrid component is replaced with its matching “empty” vector: pBR-α (pPrey-empty), pRM16 (pAdapter-empty), or pSS1 (pBait-empty). β-galactosidase activity is shown in Miller units, and the dashed horizontal line represents the basal activity level defined by the highest negative-control value. Values significantly above this line indicate a positive interaction. Data are shown as the mean of three biological replicates and error bars indicate one standard deviation from the mean. (b) Normalized data from panel (a) showing the McaS-Hfq B3H interaction plotted as the fold-stimulation of *lacZ* activity, where the β-galactosidase activity of the sample containing all three hybrid components has been divided by the highest negative-control value. We rarely find large changes in the negative control values between constructs and typically show B3H data in this normalized manner when comparing across multiple interactions. In this example, the fold-stimulation was calculated as (187±9)/(33±3) = 5.7±0.6 (mean±error). The propagated error of this fold-stimulation ratio was calculated as 5.7x[(9/187)^2^+(3/33)^2^]^1/2^ = 0.6 (see Methods 3.4.10 for the error propagation formula). B3H data for these constructs were first reported in reference [12] using a different pAdapter plasmid. **Alternative Text for Figure 4:** This figure contains a set of two bar graphs illustrating the quantitative results of a liquid beta-galactosidase assay measuring the interaction between the Hfq protein and the McaS sRNA. Absolute beta-galactosidase activity is shown on the left in panel A. The vertical Y-axis shows beta-galactosidase activity in Miller units, with a scale from 0 to 200. The horizontal X-axis displays four distinct plasmid combinations. The first bar is dark grey and represents the full tripartite complex containing all three hybrid components: pPrey-Hfq, pBait-McaS and pAdapter. This bar shows a robust signal of 187 Miller units with error bars representing 9 Miller units. The following three bars are light grey and represent negative controls where one functional component is missing in each. The values of these bars are 31 for pPrey-empty, 33 for pAdapter-empty, and 19 for pBait-empty, with vertical error bars ranging in values from 2 to 5. A horizontal dashed line is shown at 33 Miller units. Panel B shows a bar plot with a Y-axis labeled fold-stimulation, with a scale from 0 to 6. Along the X-axis is a single vertical dark grey bar labeled Hfq-McaS. The bar has a value of 5.7 and vertical error bars with a value of 0.6.

### 3.4 Quantitative β-galactosidase Assay

1. Measure and record the OD_600_ of each well using a microplate reader.
2. Make lysis buffer by adding 1.2 μL rLysozyme (400U/µL) to 1.2 mL PopCulture Reagent, providing enough for 1 96-well plate. Make this mixture fresh for each experiment.
3. Add 10 µL of lysis buffer (PopCulture + rLysozyme) per well to a fresh 96-well plate. For this step and those following, use a multichannel pipette.
4. Once cells in individual wells reach mid-log phase (see note 17), transfer 100 µL of cells to each well containing 10 µL lysis buffer. Pipette up and down to mix well and incubate at room temperature for 30 minutes.
5. Dilute 4 mL ONPG (4 mg/mL) and 43.2 µL BME into 16 mL Z buffer. Add 150 µL of this freshly prepared working assay substrate into each well of a fresh 96-well plate (see note 7).
6. Add 30 µL of lysed cells to each well of working assay substrate. In order to avoid air bubbles, do not fully depress the pipette or mix further.
7. Immediately place the plate in a microplate spectrophotometer and measure the OD_420_ at 28°. Collect data every 60 seconds for 60 minutes.
8. Normalize the slope of the OD_420_ data by OD_600_ measurements to obtain β-galactosidase activity in Miller units using the formula:

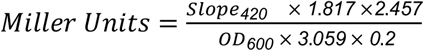

where 1.817 and 2.457 are constants, 3.059 is the pathlength adjustment, and 0.2 is the volume of cells used in the assay.
9. Data can be expressed as “fold-stimulation” of *lacZ* activity by dividing the experimental Miller Units by the highest value obtained from the three negative controls (Fig. 4).
10. When calculating fold-stimulation values, the error associated with each of the mean values (‘a’ and ‘b’) in the fold-stimulation ratio can be propagated using the following equation:

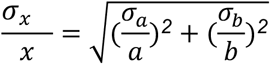

where x is the fold-stimulation value derived from the ratio of a and b, and σ_a_, σ_b_, and σ_x_ represent the standard deviation of a and b, and the propagated error of x, respectively.

## 4. Notes

1. For Hfq-RNA interactions, the B3H system can detect interactions with dissociation constants (K_D_) up to between 25 and 100 nM. However, this threshold likely varies depending on the specific RNA-protein pair; for example, the threshold for detection was a K_D_ value of approximately 25 nM for DsrA-Hfq interactions and in the 100 nM range for poly(A)-Hfq interactions (Wang et al. 2021). Additionally, plate-based assays can show higher sensitivity than liquid assays for interactions near the detection limit (see note 8).
2. The deletion of *hfq* in the reporter strain KB473 introduces a modest growth defect relative to parent FW102 -62-O_L_2 cells. If transformed cells grow too slowly for practical use (*i*.*e*. subcultures taking longer than two and a half hours to reach mid-log), the assay can be attempted in an *hfq*^*+*^ strain, though be aware that endogenous Hfq may compete with the prey protein for certain bait RNAs, as has been observed for ProQ-RNA interactions [10, 15].
3. In this protocol, pCW17 is the primary adapter plasmid used to express the CI-MS2^CP^ fusion protein under the control of a constitutive promoter. The corresponding negative control is pRM16, which shares the same constitutive promoter as pCW17 but lacks the MS2^CP^ moiety (encoding only the λCI protein). While pRM16 is the best matched control for pCW17, the original λCI-empty vector, pAC-λCI (Addgene #53730), which utilizes an IPTG-inducible *lacUV5* promoter, can also be used for simplicity. In practice, baseline β-galactosidase activity from both plasmids is very similar at the IPTG concentrations used.
4. For RNA baits that encode their own transcription terminator (*e*.*g*. bacterial sRNAs or 3’UTRs), a simple pBait vector may be used (e.g. pCH1 or the GC-clamped pSS1. For bait RNAs lacking an intrinsic terminator (*e*.*g*. 5’UTRs or fragments of open reading frames), a pBait vector providing a *trpA* terminator should be used (e.g. pHL6 or the GC-clamped pSS2). See Figure 2 for schematics of these pBait scaffolds. For more information about GC clamps, see note 5.
5. The hybrid RNA component of the B3H assay must fold correctly to ensure the MS2^hp^ is accessible to the adapter and the bait RNA is available for the prey protein. The initial B3H used pCH1 (Addgene #174663) and pHL6 as the primary bait vectors [10]. While the original vectors are functional, the updated constructs pSS1 and pSS2 are now recommended as the primary starting points for cloning new bait constructs. The optimized vectors feature 5- and 7-bp GC clamps flanking the MS2^hp^ or insertion restriction sites, respectively, to isolate the target RNA’s secondary structure from the MS2^hp^ and/or *trpA* terminator. See Figure 2 for schematics of these pBait scaffolds. These GC clamps — first used in the yeast-three hybrid system [16,17] — are computationally predicted to facilitate proper RNA folding, and improve the fold-stimulation values of several Hfq-RNA B3H interactions [12]. Since misfolding of either the MS2^hp^ or the RNA bait is a potential cause of false-negative results in the system, it is recommended to computationally predict the secondary structure of the hybrid RNA (*e*.*g*. with the “forna” web server (Kerpedjiev et al. 2015)) in order to minimize misfolding.
6. Examples of (F)orward and (R)everse primers to amplify inserts for vector ligation. The specific sequence of the 5’ “landing pad” on each primer is not critical, but incorporating at least four additional bases outside of the restriction site ensures that restriction enzymes can bind efficiently to PCR products for digestion. pPrey F Hfq: 5’-ATAAGAAT GCGGCCGC A GCTAAGGGGCAATCTTTACAAGATCC-3’ (landing pad - NotI site - extra ‘A’ - complementarity to 5’ end of DNA encoding prey protein) pPrey R Hfq: 5’ CTTC GGATCC TTATTCGGTTTCTTCGCTGTCCTG-3’ (landing pad - BamHI site - complementarity to 3’ end of DNA encoding prey protein) pBait F McaS: 5’-TCCC CCCGGG ACCGGCGCAGAGGAG-3’ (landing pad - XmaI site - complementarity to 5’ end of DNA encoding bait RNA) pBait R McaS: 5’-CCGGCC AAGCTT AAAAAATAGAGTCTGTCGACATCCGCC=3’ (landing pad - HindIII site - complementarity to 3’ end of DNA encoding bait RNA)
7. BME is essential for keeping the β-galactosidase enzyme in a reduced, active state. It must be added to the Z-buffer freshly for each assay. Additionally, to prevent hydrolysis, ONPG stocks should be stored at -20°C and thawed just before use.
8. While the primary focus of this protocol is the quantitative liquid assay, results can also be visualized with a complementary qualitative readout using X-gal indicator plates. Plate-based assays can provide increased sensitivity and consistency for some low-signal interactions. For more information, refer to the procedure in reference [2].
9. In addition to the analysis of specific protein-RNA interactions covered by this protocol, the B3H system is easily adapted for forward genetic mutagenesis screens. For detailed instructions on executing mutagenesis screens, refer to reference [2].
10. The longest bait RNA that we have successfully used in B3H assays is ∼230 nt [12], though most are between 100 and 150 nts. While longer RNAs may be viable in the system, it is recommended to use the smallest RNA fragment that will support the interaction of interest.
11. When preparing vectors for cloning, it can be easier to digest vectors that contain inserts (*e*.*g*. pKB817, pLN34, pLN43) rather than the corresponding empty vectors that serve as negative controls. The presence of an insert ensures sufficient space between the restriction sites to efficiently cleave the DNA, and digestion by both enzymes can be easily verified by the appearance of the shorter insert product on an agarose gel.
12. Any standard preparation of competent cells should be acceptable. We typically use chemically competent cells made by resuspending cells into a buffer made with 15% (v/v) glycerol, 10 mM MnCl_2_, 50 mM CaCl_2_, and 10 mM MES pH 6.3. For additional details, see reference [2]. For addressing low transformation efficiencies, see note 13.
13. Simultaneous transformation of three plasmids can result in low colony counts. If transformation efficiency is insufficient, pre-transform the reporter strain with the pAdapter plasmid and prepare competent cells that already contain this plasmid. This allows for a more efficient double transformation of the pBait and pPrey plasmids. While this approach does not allow for the pAdapter (λCI)-empty negative control, the use of pPrey and pBait empty negative controls still ensures that B3H signal depends on both of the unique elements of each detected interaction.
14. When conducting large-scale experiments, it can be advantageous to perform transformations directly in 96-well plates without first isolating single colonies. For a detailed protocol, refer to reference [2].
15. We have found that different B3H interactions are optimized with different IPTG concentrations and recommend testing a concentration range such as 0, 10, 25 and 50 μM in subcultures. The presence of IPTG in overnight cultures has been found to be helpful for Hfq-RNA interactions [11], but was not used in B3H assays with either ProQ or Khp proteins [10, 13]. Slow growth of bacteria in the presence of IPTG can indicate toxicity of the prey fusion protein. In such cases, reduce IPTG concentrations (e.g., to 0–10 μM) to alleviate the fitness cost while maintaining enough protein for detection. In particular, we have found that omission of IPTG from overnight cultures can help avoid inconsistent cell growth.
16. Inconsistent OD_600_ readings across a 96-well plate are often caused by media evaporation. Ensure plates are covered during all incubation steps. If using an incubator without humidity control, place an uncovered container of warm water at the base to maintain 70–80% humidity.
17. Bacterial cells grown in 96-well formats often reach mid-log phase at different rates due to variations in growth rates or the cell density of overnight cultures. To ensure consistency, it is recommended to monitor the OD_600_ frequently (*e*.*g*. every 20 minutes after an initial growth period of 60-90 minutes) and lyse “fast-growers” immediately as they hit mid-log, allowing slower wells to continue growing. Lysed samples can remain at room temperature for up to 3 hours without significant loss of β-galactosidase activity.
18. To facilitate the study of bacterial 5’UTR and ORF fragments, pBait vectors pHL6 and pSS2 encode a UAA stop codon downstream of the HindIII site. This allows any initiating ribosomes to terminate translation and dissociate from the bait RNA without stalling at its 3’ end. If a bait RNA does not contain a bacterial ribosomal binding site and start codon, this stop codon may be disregarded. For mRNA fragments containing these elements, primers should be designed to ensure that the UAA stop codon remains in frame with the mRNA’s reading frame.

## 6. Acknowledgements

Recent work in our laboratory developing this bacterial three-hybrid assay has been supported by funding from National Institutes of Health [2R15GM135878 to K.E.B.], the Camille and Henry Dreyfus Foundation, and Mount Holyoke College.

## References

1. Berry KE, Hochschild A (2018) A bacterial three-hybrid assay detects Escherichia coli Hfq-sRNA interactions in vivo. Nucleic Acids Res 46: e12

2. Stockert OM, Gravel CM, Berry KE (2022) A bacterial three-hybrid assay for forward and reverse genetic analysis of RNA–protein interactions. Nat Protoc 17: 941–961

3. Dove SL, Joung JK, Hochschild A (1997) Activation of prokaryotic transcription through arbitrary protein-protein contacts. Nature 386: 627–630

4. Dove SL, Hochschild A (2004) A bacterial two-hybrid system based on transcription activation. In: Fu H (ed) Protein-Protein Interactions. Methods in Molecular Biology 261: 231–246. Humana Press, New Jersey

5. Rio DC (2014) Electrophoretic mobility shift assays for RNA–protein complexes. Cold Spring Harb Protoc doi:10.1101/pdb.prot080721

6. Hall KB, Kranz JK (1999) Nitrocellulose filter binding for determination of dissociation constants. In: Haynes SR (ed) RNA-Protein Interaction Protocols. Methods in Molecular Biology 118: 105–114. Humana Press, New Jersey

7. Andresen L, Holmqvist E (2018) CLIP-Seq in bacteria: Global recognition patterns of bacterial RNA-binding proteins. In: Carpousis AJ (ed) Methods in Enzymology 612: 127–145. Academic Press, Massachusetts

8. Li L, Förstner KU, Chao Y (2018) Computational analysis of RNA–protein interactions via deep sequencing. in Wang Y, Sun, M (eds) Transcriptome Data Analysis 1751: 171–182. Springer, New York

9. Zambelli F, Pavesi G (2015) RIP-Seq data analysis to determine RNA–protein associations. In: Picardi E (ed) RNA Bioinformatics 1269: 293–303. Springer, New York

10. Pandey S, Gravel CM, Stockert OM, Wang CD, Hegner CL, LeBlanc H, Berry KE (2020) Genetic identification of the functional surface for RNA binding by Escherichia coli ProQ. Nucleic Acids Res 48: 4507–4520

11. Wang CD, Mansky R, LeBlanc H, Gravel CM, Berry KE (2021) Optimization of a bacterial three-hybrid assay through in vivo titration of an RNA–DNA adapter protein. RNA 27: 513–526

12. Nguyen LD, LeBlanc H, Berry KE (2025) Improved constructs for bait RNA display in a bacterial three-hybrid assay. Sci Rep 15: 3820

13. Nguyen KT, Lett NW, Gravel CM, Jo S, Shi Y, Narayan M et al (2026) Molecular genetic characterization of bacterial KH-domain proteins. BioRxiv 2026.03.16.712068

14. Whipple FW (1998) Genetic analysis of prokaryotic and eukaryotic DNA-binding proteins in Escherichia coli. Nucleic Acids Res. 26, 3700–3706

15. Stein EM, Kwiatkowska J, Basczok MM, Gravel CM, Berry KE, Olejniczak M (2020) Determinants of RNA recognition by the FinO domain of the Escherichia coli ProQ protein. Nucleic Acids Res 48: 7502–7519

16. Cassiday LA, Maher LJ (2001) In vivo recognition of an RNA aptamer by its transcription factor target. Biochemistry 40: 2433–2438

17. Bernstein D (2002) Analyzing mRNA–protein complexes using a yeast three-hybrid system. Methods 26: 123–141

